# Transcriptomic Profiling of Developing Melanocortin Neurons Reveals a Role for Prdm12 in Energy Balance

**DOI:** 10.1101/753434

**Authors:** Xiameng Chen, Steven C. Wyler, Li Li, Amanda G. Arnold, Rong Wan, Pin Xu, Chen Liu

## Abstract

The hypothalamus is a critical regulator of many physiological processes essential for life. In the adult brain, each anatomically-defined hypothalamic nucleus consists of functionally heterogeneous neuronal subpopulations that dictate distinct survival behaviors. Nevertheless, how rich neuronal identities are established in the developing hypothalamus remains poorly understood.

Despite their opposing actions on food intake, POMC and NPY/AgRP neurons in the arcuate nucleus of the hypothalamus (ARH) are derived from the same progenitors that give rise to ARH neurons. However, the mechanism whereby common neuronal precursors subsequently adopt either the anorexigenic (POMC) or the orexigenic (NPY/AgRP) identity remains elusive.

We hypothesize that POMC and NPY/AgRP cell fates are specified and maintained by distinct intrinsic factors. In search of them, we profiled the transcriptomes of developing POMC and NPY/AgRP neurons with whole-genome RNA sequencing (RNA-seq). Moreover, cell-type-specific transcriptomic analyses revealed transcription regulators that are selectively enriched in either population, but whose developmental functions are unknown in these neurons.

Among them, we found the expression of the PR domain-containing factor 12 (*Prdm12*) was enriched in POMC neurons but was absent in NPY/AgRP neurons. Selective ablation of *Prdm12* in postmitotic POMC neurons led to early-onset obesity, accelerated linear growth, as well as impaired glucose tolerance, which recapitulates symptoms of POMC/MC4R deficiency in humans. These findings, therefore, establish a previously-unrecognized role for *Prdm12* in energy balance. Furthermore, the combination of cell-type-specific genomic and genetic analyses provides a means to dissect cellular and functional diversity in the developing hypothalamus as well as the developmental origins of diverse survival behaviors.

---

Serving as the interface between the nervous and endocrine systems, the hypothalamus is a critical regulator of many physiological processes essential for life. Recent advances in optogenetic, chemogenetic, and imaging techniques have enabled the isolation of individual hypothalamic circuits that control satiety ^1^, thirst ^2^, aggression ^3^, and many more. These studies further demonstrate that each anatomically-defined hypothalamic nucleus consists of functionally heterogeneous neuronal subtypes that dictate distinct survival behaviors. On the other hand, how rich neuronal identities are established in the developing hypothalamus remains poorly understood ^4^.

Within the ARH, earlier studies have uncovered a transcriptional cascade that sequentially specifies postmitotic neuronal precursors ^5-8^. However, the mechanism whereby common neuronal precursors subsequently adopt either the anorexigenic (POMC) or the orexigenic (NPY/AgRP) identity remains elusive. While fate specification has been thoroughly investigated in other parts of the nervous system, it remains an understudied subject in the hypothalamus. For example, the dynamic expression patterns of peptidergic markers make it difficult to track POMC and NPY/AgRP neurons during development ^9^. Moreover, since cell bodies of these neurons are closely intermingled with each other, existing databases on gene expression (e.g. Genepaint and Allen Brain Library) offer few clues for cell-type-specific transcripts behind fate choices. To overcome these hurdles, we purified genetically-labeled POMC and NPY/AgRP from the developing mouse hypothalamus and profiled their transcriptomes using whole-genome RNA sequencing.

To track developing POMC and NPY/AgRP neurons, we took advantage of two well-characterized transgenic mice (POMC^eGFP^ and NPY^hrGFP^) in which POMC and NPY/AgRP neurons are genetically labeled by the expression of green fluorescent proteins ^10,11^. Despite a shared birthdate ^9^, POMC and NPY/AgRP neurons launch their peptidergic marker expression at distinct developmental periods. *Pomc* mRNA first appears in the hypothalamic ventricular zone at embryonic (E) day E10.5 ^6^. It is transiently expressed in other neurons before being committed to prospective POMC neurons after E14.5 ^12^. In comparison, *Npy* mRNA is not present in the presumptive ARH until E13.5 ^9^. Consistent with endogenous *Pomc* and *Npy*, we observed the onset of POMC-eGFP and NPY-hrGFP expression at E10.5 and E13.5 respectively in the developing ARH (Supplementary Fig. 1a-b). Although NPY/AgRP neurons co-express NPY and AgRP in the adult brain ^13^, it has been reported that the onset of *Agrp* expression within NPY/AgRP neurons did not occur until perinatally ^14^. Using AgRP-iresCre activity as a surrogate for endogenous *Agrp* expression ^15^, we detected the expression of Cre-activated tomato reporter (*Rosa26*^*Ai14*^) in only a few ARH neurons as early as E15.5 (Supplementary Fig. 1c). The number of tomato positive neurons gradually increases over time, and by postnatal day 7 (P7), reporter expression is found only in a subset of NPY-hrGFP neurons (Supplementary Fig. 1d).

Therefore, our finding supports the notion that the acquisition of the AgRP identity spans an extended postnatal period.

To uncover the intrinsic factors underlying the anorexigenic vs. the orexigenic identity, we profiled the transcriptomes of developing POMC and NPY/AgRP neurons at E15.5 while they were acquiring their adult neurotransmitter identities. Under a fluorescent dissecting microscope, we dissociated the ARH from POMC^eGFP^ or NPY^hrGFP^ embryos and purified GFP positive neurons through fluorescence-activated cell sorting (FACS, Fig. 1a). We collected 3-4 biological replicates per genotype. Each biological replicate was defined as an independent sort from pooled embryos from different litters. We extracted total RNAs from approximately five-thousand to ten-thousand neurons per replicate and subjected them to whole-transcriptome RNA sequencing (Illumina HiSeq™ 2500). We obtained an average of 77.8 million mapped reads/sample. Reads were aligned to the mouse reference genome GRCm38/mm10 assembly using HISAT2 ^16^. From there, we detected 10317 genes present in at least one of the two populations with a minimum expression of 10 transcripts per million mapped reads (TPM, Supplementary Table 1).

**Fig. 1:**
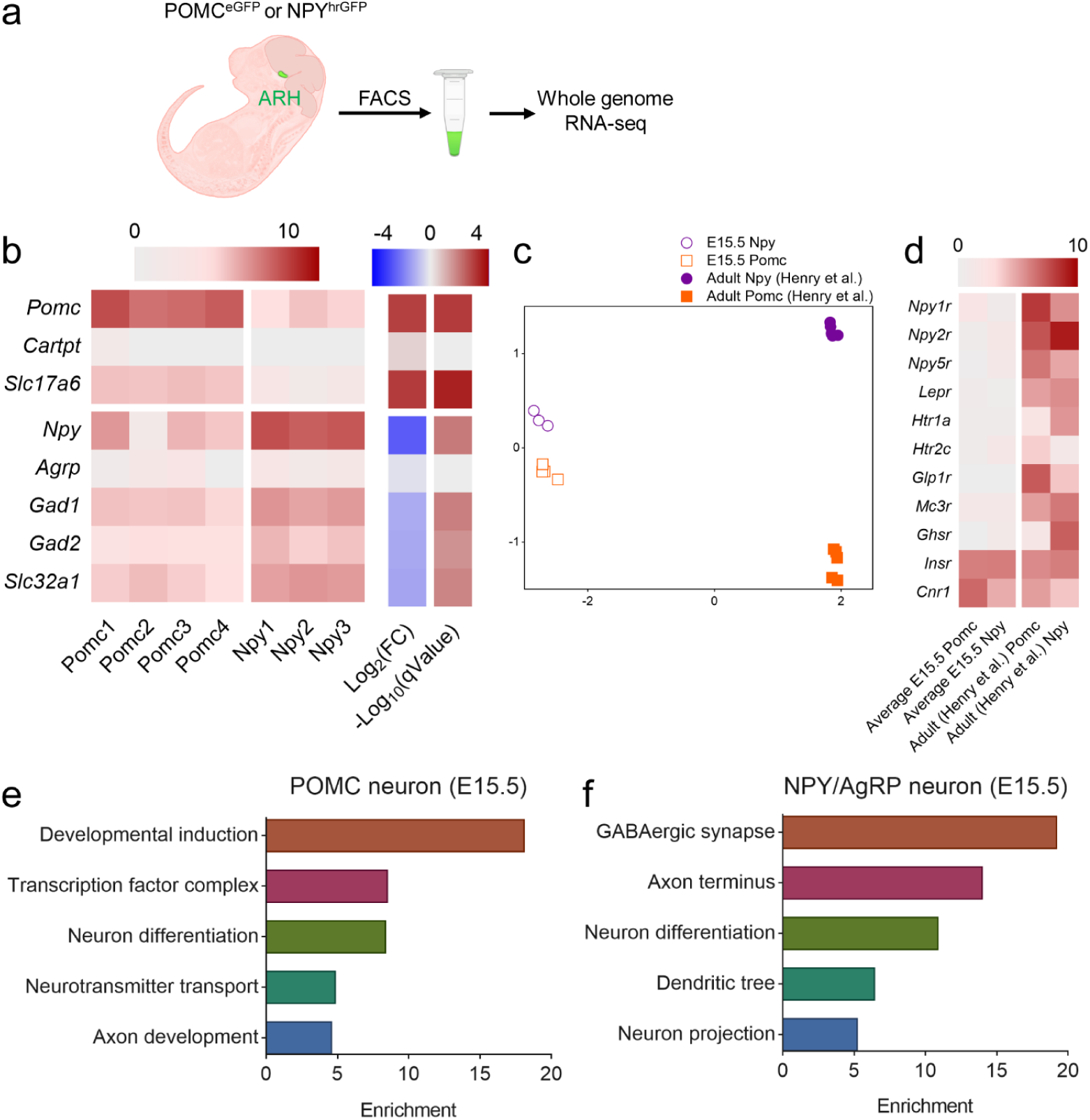
Transcriptomic profiling of embryonic POMC and NPY/AgRP neurons. **a**, Schematic of isolation and profiling transcriptomes of E15.5 POMC and NPY/AgRP neurons. **b**, Heatmap of cell fate marker expression in POMC and NPY/AgRP neurons. Rows are replicates for each gene. Log2(FC), log fold change of gene expression in POMC neurons over that in NPY/AgRP neurons. q-Value: adjusted p-value; FDR ≤ 5% (Benjamin-Hochberg correction). **c**, Multidimensional scaling (MDS) projection of distance (1-correlation coefficient) of E15.5 POMC and NPY/AgRP neurons compared to data extracted from previously published adult (fed only) POMC and NPY/AgRP neurons (Henry et al.). **d**, Heat map of the average of replicates for embryonic (columns 1 and 2) and adult (columns 3 and 4, Henry et al.) expression. Scale: log2(TPM+1). **e, f** Gene ontology for enriched genes in POMC or NPY/AgRP neurons respectively.

Our analysis identified 120 differentially expressed genes (DEGs) between POMC and NPY/AgRP neurons at E15.5 (Supplementary Table 2 and see ‘Materials and methods’ for criteria). As expected, transcripts for *Pomc* and *Npy* were highly enriched in POMC and NPY/AgRP neurons respectively (Fig. 1b). In contrast, *Cart* and *Agrp* whose expression comes on late during development were barely detectable at this stage. Other than the peptidergic markers, we found an enrichment of *Slc17a6* (vGLUT2), a marker for glutamatergic neurons in POMC neurons, whereas markers for GABAergic neurons such as *Slc32a1* (vGAT), *Gad1* and *Gad2* were over-represented in NPY/AgRP neurons (Fig. 1b). Using similar approaches, Henry et al. profiled the transcriptomes of adult POMC and NPY/AgRP neurons ^17^. We compared the embryonic and adult datasets using multidimensional scaling analyses and found a clear separation in gene expression patterns of the same neurons (Fig. 1c). For example, adult POMC and NPY/AgRP express receptors for multiple metabolic hormones such as leptin, insulin, ghrelin, and serotonin. However, most of these receptors—except for the insulin receptor and the cannabinoid receptor 1—were expressed at very low levels (between 0 and 6.3 TPM) at E15.5 (Fig. 1d). Along the same line, pathway analyses revealed an enrichment of genes involved in G-protein coupled receptor signaling, endoplasmic reticulum stress and circadian rhythm in adult POMC and NPY/AgRP neurons ^17^. In contrast, those involved in neuronal growth, differentiation, and axon development (Fig. 1e-f, Supplementary Table 3) were overrepresented at E15.5. Collectively, these findings demonstrate that developing POMC and NPY/AgRP neurons possess distinct transcription profiles compared to their adult counterparts.

We focused our analyses on transcription factors (TFs) as they play a critical role in neuronal differentiation and fate specification. We detected transcripts of 1106 TFs in either POMC or NPY/AgRP neurons at E15.5 (Supplementary Table 4). While the vast majority of these factors were present at equivalent levels in both neurons, 29 of them showed differential enrichment in one of the two groups (Fig. 2a). As expected, the basic helix-loop-helix factor *Nhlh2* is enriched in POMC neurons and controls the levels of prohormone convertase I (*Pcsk1)*, the enzyme that converts POMC to α-MSH ^18^. Moreover, TFs such as *Nkx2*.*1, Tbx3, Dlx1*, and *Dlx2* are initially expressed in a broad domain of progenitors during early differentiation ^5,19,20^. We found that their expression persists in postmitotic neurons and are selectively enriched in either POMC (*Nkx2*.*1* and *Tbx3*) or NPY/AgRP (*Dlx1* and *Dlx2*) neurons (Fig. 2a-c). These findings are consistent with recent reports that these factors continue to play a role in POMC or NPY/AgRP neurons after neurogenesis ^21-23^.

**Fig. 2:**
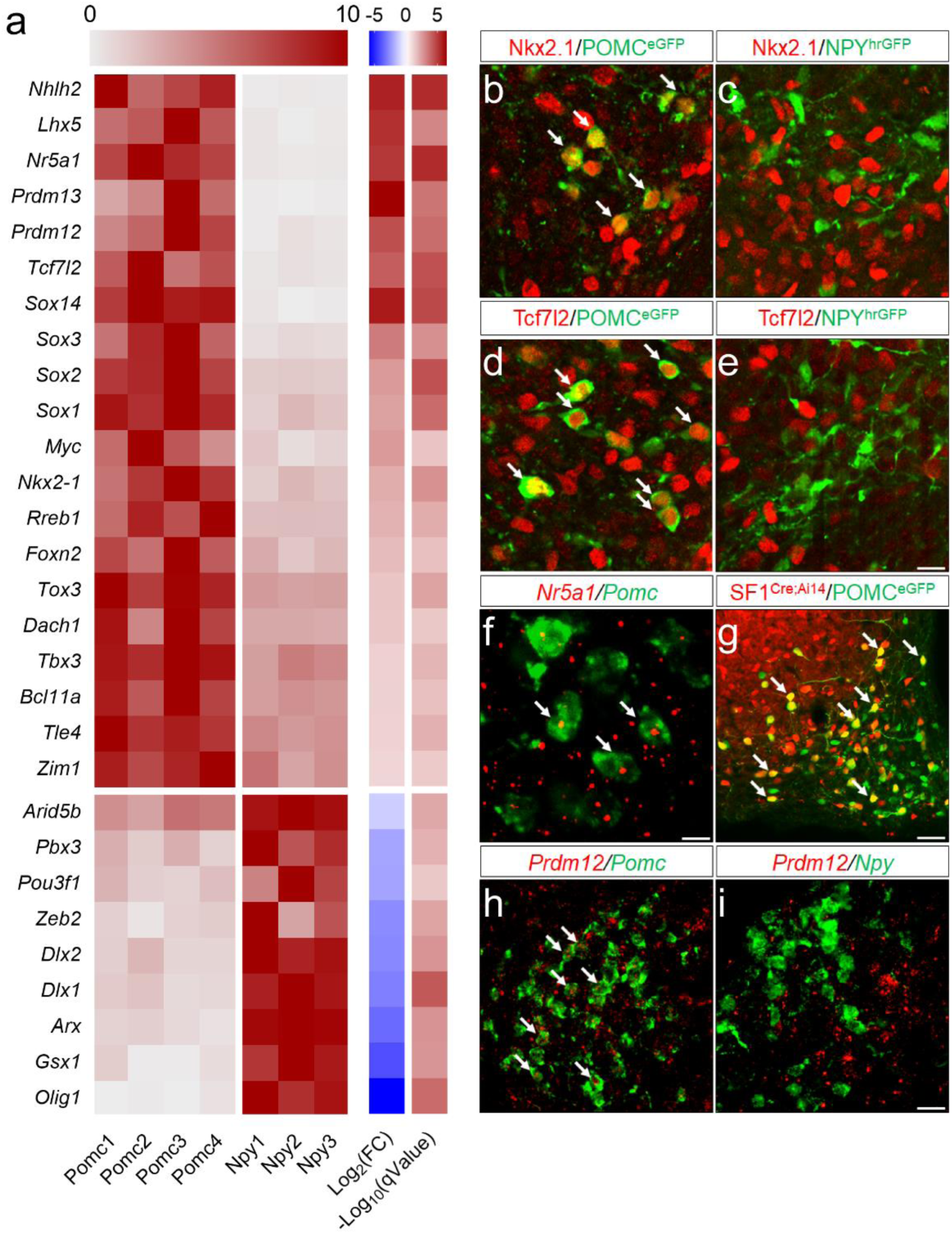
TFs selectively enriched in either POMC or NPY/AgRP neurons at E15.5. **a**, Heatmap of 29 differently expressed TFs at E15.5. Each row is normalized to the maximum expression value for each replicate sample. Log2(FC): log fold change of gene expression in POMC neurons over that in NPY/AgRP neurons. q-Value: adjusted p-value; FDR ≤ 5% (Benjamin-Hochberg correction). **b-e**, Double-staining of Nkx2.1 (red), Tcf7l2 (red), and GFP (green) in the ARH at E15.5. **f**, Double fluorescent *in-situ* hybridization (RNAscope) of *Nr5a1* mRNA (red) and *Pomc* mRNA (green) at E15.5. **g**, Immunofluorescence of tomato (red) and GFP (green) in the adult ARH. **h, j**, Double fluorescent *in-situ* hybridization (RNAscope) of *Prdm12* mRNA (red), *Pomc* mRNA (green), or *Npy* mRNA (green) in the ARH at E15.5. Arrowheads indicate neurons that express both epitopes or mRNAs. Scale bar is 5 µm in **f**, 10 µm in (**b-e**), 20 µm in (**h, i**), and 50 µm in **g**.

In addition to the TFs that have an established role, our analyses identified many TFs that have yet been implicated in the development of ARH neurons. Particularly, the establishment and maintenance of the anorexigenic POMC neuron identity is highly relevant to childhood obesity as mutations in *Pomc* cause severe early-onset obesity in humans ^24^. However, genetic factors that regulate *Pomc* expression remain largely unknown. We found several members of the Sox proteins (*Sox1, Sox2, Sox3*, and *Sox14*) were enriched in POMC neurons (Fig. 2a). *de Souza et al*. previously characterized upstream enhancer elements (nPE1 and nPE2) that were both necessary and sufficient to drive reporter gene expression in hypothalamic POMC neurons ^25^. Analyses of these sequences revealed several conserved Sox binding sites across multiple mammalian species (Supplementary Fig. 2). These findings, therefore, raise the possibility that Sox proteins may directly interact with upstream enhancer sequences to regulate *Pomc* transcription. Moreover, we found that developing POMC neurons, but not NPY/AgRP neurons, express mRNAs and proteins of the transcription factor 7-like 2 (*Tcf7l2*, Fig2. a, d, e). Polymorphisms of *Tcf7l2* are among the most significant genetic markers associated with type 2 diabetes across multiple ethnic groups ^26^. Previous studies largely focused on its role in peripheral organs such as the pancreatic islets and the liver ^27^. However, given the importance of POMC neurons in coordinating glucose metabolism ^28^, it is possible that these neurons may serve as a critical node where *Tcf7l2* programs glucose homeostasis at critical developmental periods. Furthermore, the nuclear receptor *Nr5a1* (also known as Steroidogenic factor 1, SF1) has been known as a selective marker for VMH neurons during development ^29^. Its expression within the developing ARH, however, is controversial. While an earlier study reported Nr5a1 proteins in developing POMC neurons ^6^, a subsequent analysis failed to corroborate this finding ^30^. Using cell-type-specific RNA-seq, we detected *Nr5a1* mRNAs in POMC neurons (Fig. 2a, f). In support of this finding, fate-mapping of *SF1-Cre* ^31^ expressing neurons revealed that a subset of adult POMC neurons (37.6% ± 5.5%) expresses the Cre-activated tomato reporter (Fig. 2g). Since *Nr5a1* is absent from adult POMC neurons^17^, our finding suggests that *Nr5a1* is transiently expressed in POMC neurons during development. However, a developmental role for *Nr5a1* within POMC neurons remains to be determined. Furthermore, whether its expression demarcates a functionally-unique subset of POMC neurons warrants further investigations. Finally, our analyses uncovered an enrichment of two PR-domain containing factors *Prdm12* and *Prdm13* in developing POMC neurons (Fig. 2a, h, i). Both factors play an important role in lineage specification in the developing spinal cord ^32-34^. However, whether these factors play a similar role in the hypothalamus has not been studied.

To test whether POMC-enriched TFs are necessary for the anorexigenic neuron identity, we turned to mouse genetics to interrogate their functions *in vivo*. We first targeted *Prdm12* since it has not been implicated in the development of hypothalamic neurons nor in energy balance. Prior to the first appearance of *Pomc*, we detected *Prdm12* mRNA in the ventral hypothalamic neuroepithelium at E9.5 (data not shown). Moreover, it is present within POMC neurons when they first emerge at E10.5 (Fig. 3a). To investigate the physiological role of *Prdm12 in vivo*, we developed a conditional *Prdm12* allele (*Prdm12*^*flox*^) by inserting two *loxP* sequences flanking the endogenous exon that encodes the DNA-binding domain (Fig. 3b). Breeding of heterozygous *Prdm12*^*fl/+*^ mice produced *Prdm12*^*+/+*^, *Prdm12*^*fl/+*^, and *Prdm12*^*fl/fl*^ offspring (Fig. 3c) with the expected Mendelian ratio. The floxed exon could be removed in the presence of *Cre* recombinase resulting in a deleted *Prdm12* allele (*Prdm12*^*Δ*^, Fig. 3b, d). Meanwhile, quantitative PCR (qPCR) analyses detected comparable levels of *Prdm12* mRNAs within the hypothalamus between *Prdm12*^*+/+*^ and *Prdm12*^*fl/fl*^ littermates, suggesting that the introduced *loxP* sites did not perturb endogenous *Prdm12* transcription (Fig. 3e). We bred *POMC-Cre* mice ^35^ with *Prdm12*^*fl/fl*^ mice to generate *POMC-Cre; Prdm12*^*fl/fl*^ mice in which *Prdm12* was selectively deleted in embryonic *POMC* neurons (designated hereafter as *Prdm12*^*POMCeKO*^ mice). A Cre-dependent tdTomato reporter (*Rosa26*^*Ai14*^) was used to mark *POMC-Cre* expressing neurons in *Prdm12*^*POMCeKO*^ mice as well as those in mice heterozygous for the floxed *Prdm12* allele (*POMC-Cre; Prdm12*^*fl/+*^, Fig. 3f, g). We verified the loss of *Prdm12* mRNAs in FACS-sorted tdTomato+ neurons in *Prdm12*^*POMCeKO*^ mice (Fig. 3h). Cell counting analyses found similar numbers of tdTomato+ cells present in the adult brains of both genotypes, suggesting that *Prdm12* is dispensable for the survival of POMC neurons (Fig. 3i). Remarkably, *in situ* hybridization uncovered a significant loss of *Pomc* mRNAs in the ARH of *Prdm12*^*POMCeKO*^ mice (Fig. 3j, k). In contrast, the expression of *Npy* and *Agrp* remained unaffected (Fig. 3l-3o). Importantly, immunohistochemistry for beta-endorphin, a proteolytic product of POMC, suggested that *Pomc* expression is not altered in the pituitary gland (Fig. 3p-q). Consistent with these findings, qPCR analyses confirmed a selective loss of *Pomc* transcripts among several genes known to be expressed in the ARH. Collectively, these findings suggested a hypothalamic *Pomc* deficiency in *Prdm12*^*POMCeKO*^ mice (Fig. 3r).

**Fig. 3:**
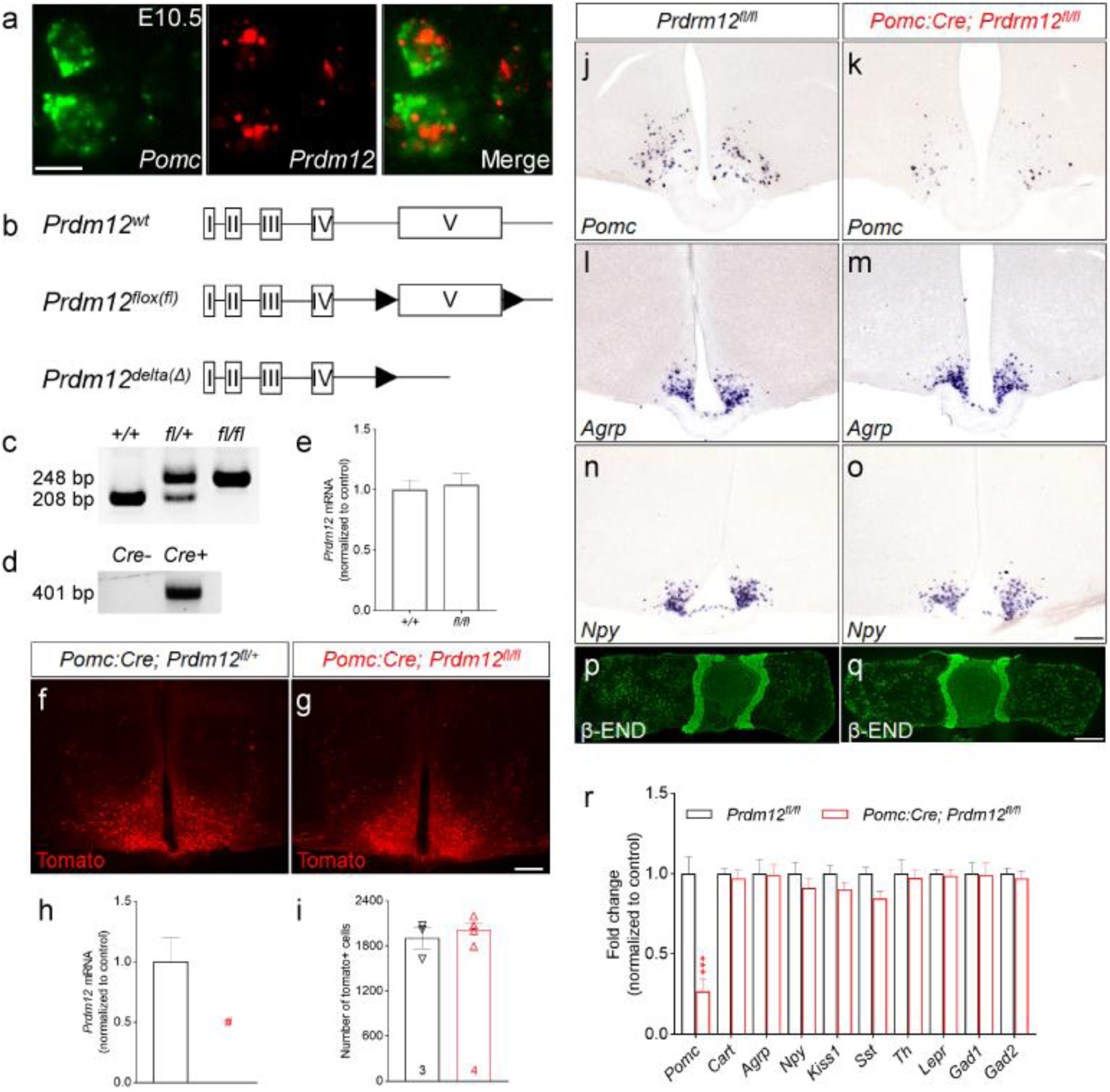
*Prdm12* is required for *Pomc* expression during development. **a**, Fluorescent *in-situ* hybridization for *Pomc* and *Prdm12* mRNAs at E10.5. **b**, Schematics of the wildtype (wt), floxed (fl), and delta(Δ) *Prdm12* alleles. **c**, PCR genotyping of *Prdm12*^*+/+*^, *Prdm12*^*fl/+*^, and *Prdm12*^*fl/fl*^ mice. **d**, Detection of the *Prdm12*^*Δ*^ allele by PCR. **e**, qPCR analysis of *Prdm12* mRNA in the hypothalamus, n=5-7. **f, g**, Immunofluorescence of tomato. **h**, qPCR analysis of *Prdm12* in FACS-sorted tomato positive neurons. n=3 replicates from pooled embryos, # not detectable. **i**, cell counts of tomato positive neurons, n=3-4. **j-k**, *In-situ* hybridization. **p, q**, Immunostaining for β-endorphin (END). **r**, qPCR analysis of gene expression in the hypothalamus. Scale bar is 5 µm for **a**, 200 µm for **f, g, j-o**, and 1mm for **p, q**. *** p<0.001.

POMC activates the downstream melanocortin 4 receptors (MC4R) to suppress appetite. Of note, genetic variants within the POMC/MC4R pathway are the most common obesity-causing mutations found in obese children ^36^. We investigated whether *Prdm12*^*POMCeKO*^ mice manifested similar metabolic deficits observed in humans ^37^. We found that *Prdm12*^*POMCeKO*^ mice had increased body weight at the time of weaning (Fig. 4a). The excessive weight gain was further exacerbated by subsequent feeding with a high-fat diet (HFD, 60 kcal% fat, D12492i Research Diets, Fig. 4b). Nuclear magnetic resonance imaging revealed that the elevated body weight was largely due to an increase in fat mass, along with a modest gain in lean mass (Fig. 4c). In support of this finding, H&E staining revealed significantly increased lipid accumulation in both the white adipose tissue and liver of *Prdm12*^*POMCeKO*^ mice (Supplementary Fig. 3). Moreover, consistent with an MC4R deficiency, we found that *Prdm12*^*POMCeKO*^ mice had an increased body length (nose to anus) compared with their littermate controls (Fig. 4d). To further unravel the physiological changes behind the excessive weight gain, we conducted metabolic chamber analyses using a new cohort of *Prdm12*^*POMCeKO*^ mice and their littermate controls. These mice had been maintained on a regular chow diet before entering the experiment at eight weeks of age. By that time, *Prdm12*^*POMCeKO*^ mice had increased fat mass (2.31±0.12g vs. 1.18±0.14g, p<0.001), but had comparable lean mass (20.61±0.59g vs. 20.48±0.47g, p=0.87). Mice were fed a chow diet during acclimation and the first three days of recording before being switched to an HFD for the next seven days. Under both diets, *Prdm12*^*POMCeKO*^ mice consumed more food during the dark phase of a day (Fig. 4e-f). On the other hand, heat production was similar between chow-fed *Prdm12*^*POMCeKO*^ and control mice (Fig. 4g). In contrast, heat production was significantly lower in *Prdm12*^*POMCeKO*^ mice during HFD exposure (Fig. 4h). Collectively, these results suggest that the HFD-induced exacerbation of weight gain is the result of a combination of increased energy intake and reduced expenditure. Finally, concomitant with obesity were perturbations in glucose homeostasis. For example, *Prdm12*^*POMCeKO*^ mice showed a deficit in glucose clearance during a glucose tolerance test (GTT, Fig. 4l). In control mice, plasma insulin levels elevated in response to the glucose challenge during the GTT. In contrast, insulin levels were significantly higher in *Prdm12*^*POMCeKO*^ mice at the baseline but did not increase during the GTT (Fig. 4j). The impairment in insulin sensitivity was further confirmed using an insulin tolerance test (ITT, Fig. 4k).

**Fig. 4:**
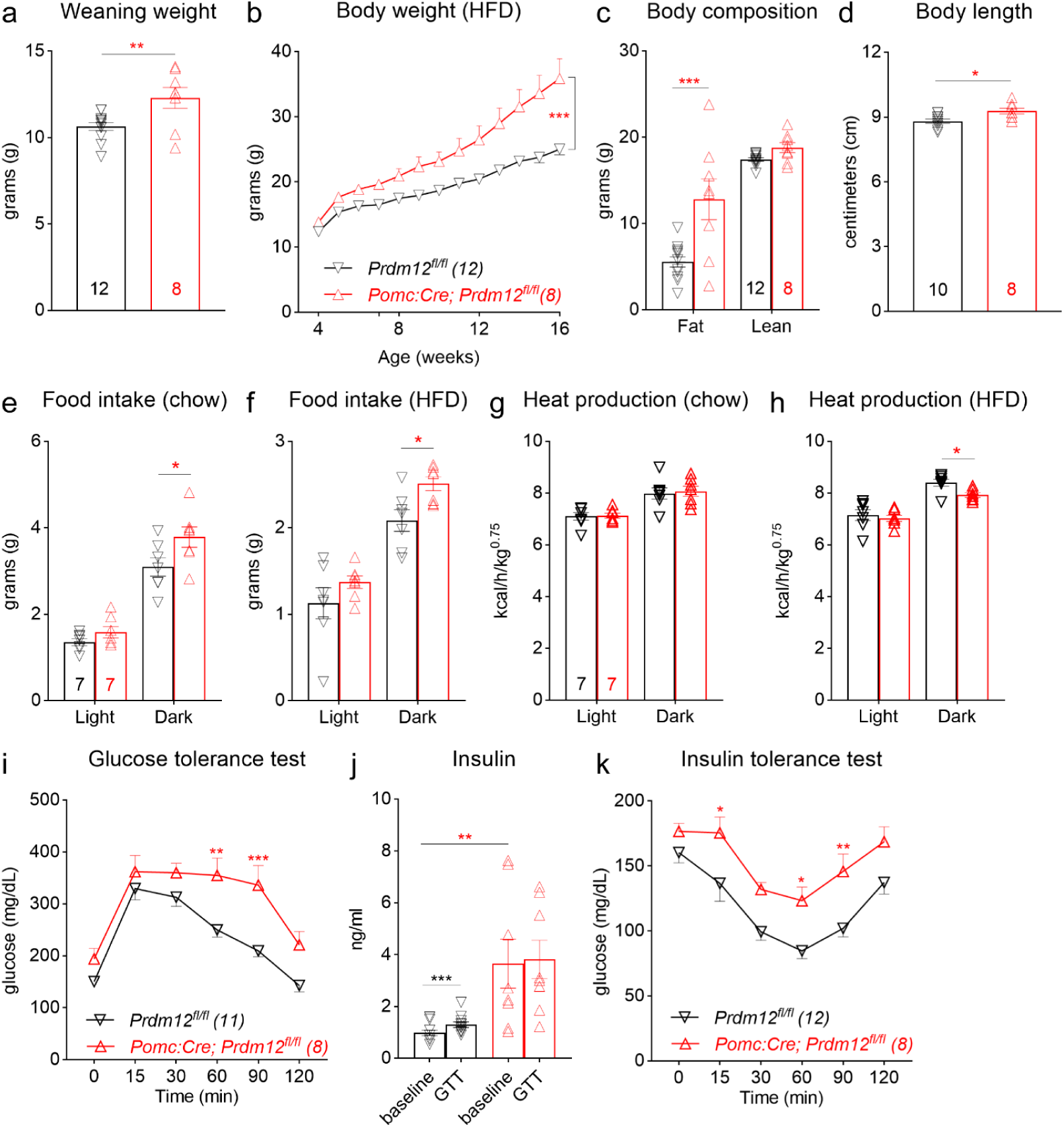
Early-onset obesity, increased linear growth, and glucose intolerance in *Prdm12*^*POMCeKO*^ mice. **a, b**, Body weight. **c**, Body composition. **d**, Body length. **e, f**, Food intake during light (12h) and dark (12h) phases of a day. **g, h**, Heat production. **i**, GTT. **j**, Serum insulin levels. **k**, ITT. * p<0.05, ** p<0.01, *** p<0.001, unpaired t-test for **a, d, j**; two-way ANOVA with Sidak’s tests for **b**,**c, e-k**.

In summary, the combination of cell-type-specific transcriptomics and mouse genetics provided a means to dissect cellular and functional diversity in the developing hypothalamus and help better understand the developmental origins of hypothalamus-controlled survival behaviors. Transcriptomic profiling of developing POMC and NPY/AgRP neurons allowed us to decipher the transcriptional codes behind the orexigenic vs. anorexigenic identity. Moreover, it offers a resource for studying other developmental milestones (e.g. axon pathfinding, synapse formation) necessary for the assembly of melanocortin feeding circuits. Finally, our studies uncovered a previously unrecognized role for *Prdm12* in *Pomc* transcription. Importantly, loss of *Prdm12* in embryonic POMC neurons led to early-onset obesity, which is characteristic of human POMC/MC4R deficiency. Therefore, the discovery of *Prdm12*, along with other POMC-enriched TFs, helps identify new genetic factors behind the rising epidemic of childhood obesity.

---

**Supplementary Figure 1.**
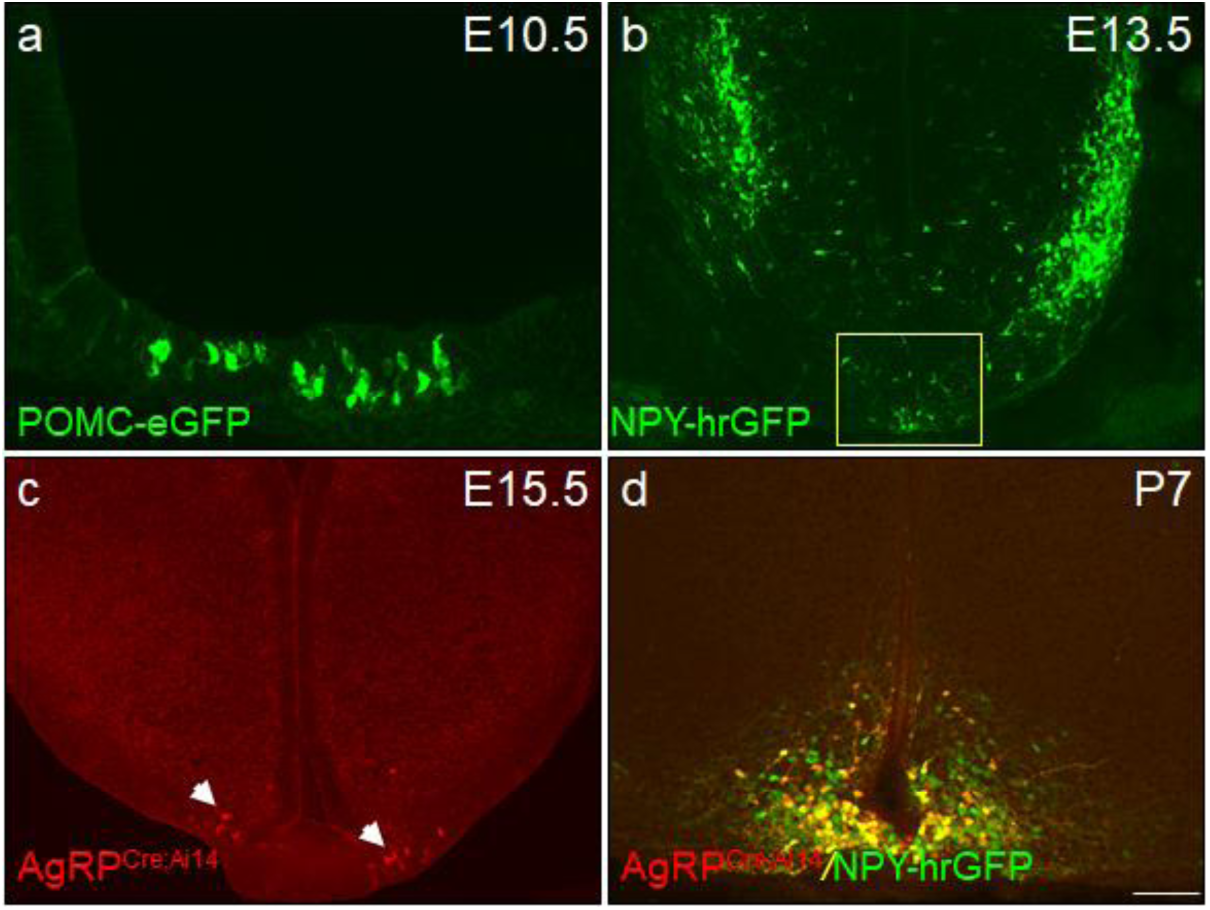
The onset of POMC-eGFP, NPY-hrGFP, and AgRP^Cre; Ai14^ in the developing ARH. **a**, Immunohistochemistry for GFP. **b**, Native fluorescence of NPY-hrGFP, yellow box, ARH. **c**, Native fluorescence of tdTomato, white arrows pointing towards tdTomato-expressing neurons. **d**, Double-labeling between tdTomato and NPY-hrGFP. Scale bar is 50 µm for **a**, and 100 µm for **b-d**.

**Supplementary Figure 2.**
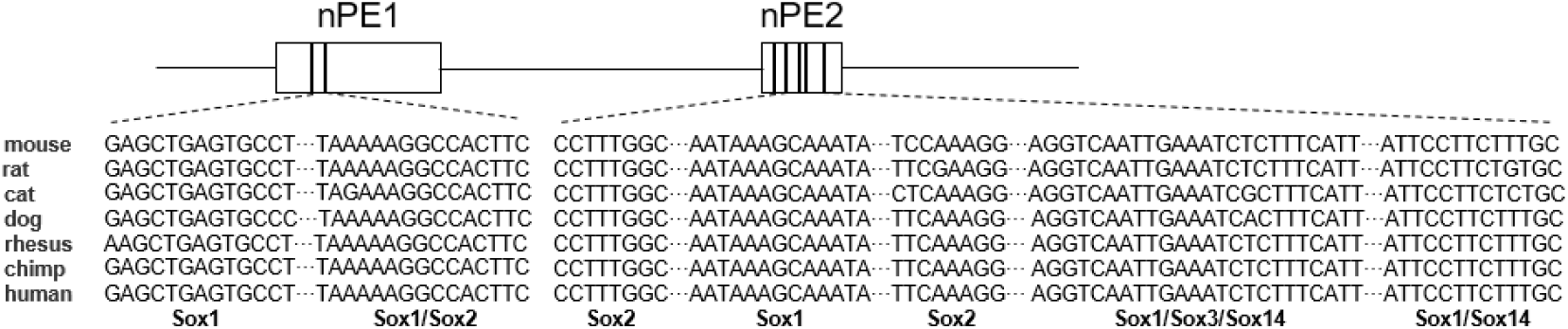
Conserved Sox binding motifs in *Pomc* neuronal enhancers nPE1 and nPE2. Sequences of mouse nPE1 (chr12:3941972-3942627, 656 bp) and nPE2 (chr12:3944600-3944917, 318 bp) defined by *de Souza* et al. were obtained using the ECR Genome Browser (https://ecrbrowser.dcode.org). Sequence alignment was conducted using the ConTra V3 web server (http://bioit2.irc.ugent.be/contra/v3/). Matrices used to predict Sox binding motifs include taipale-TCAATWNCATTGA-Sox1-DBD for Sox1, MA0143.3 for Sox2, M1592_1.02 for Sox3, and taipale-ACAATANCATTG-Sox14-DBD for Sox4 with the stringency setting (core=1.00, similarity matrix=0.95).

**Supplementary Figure 3.**
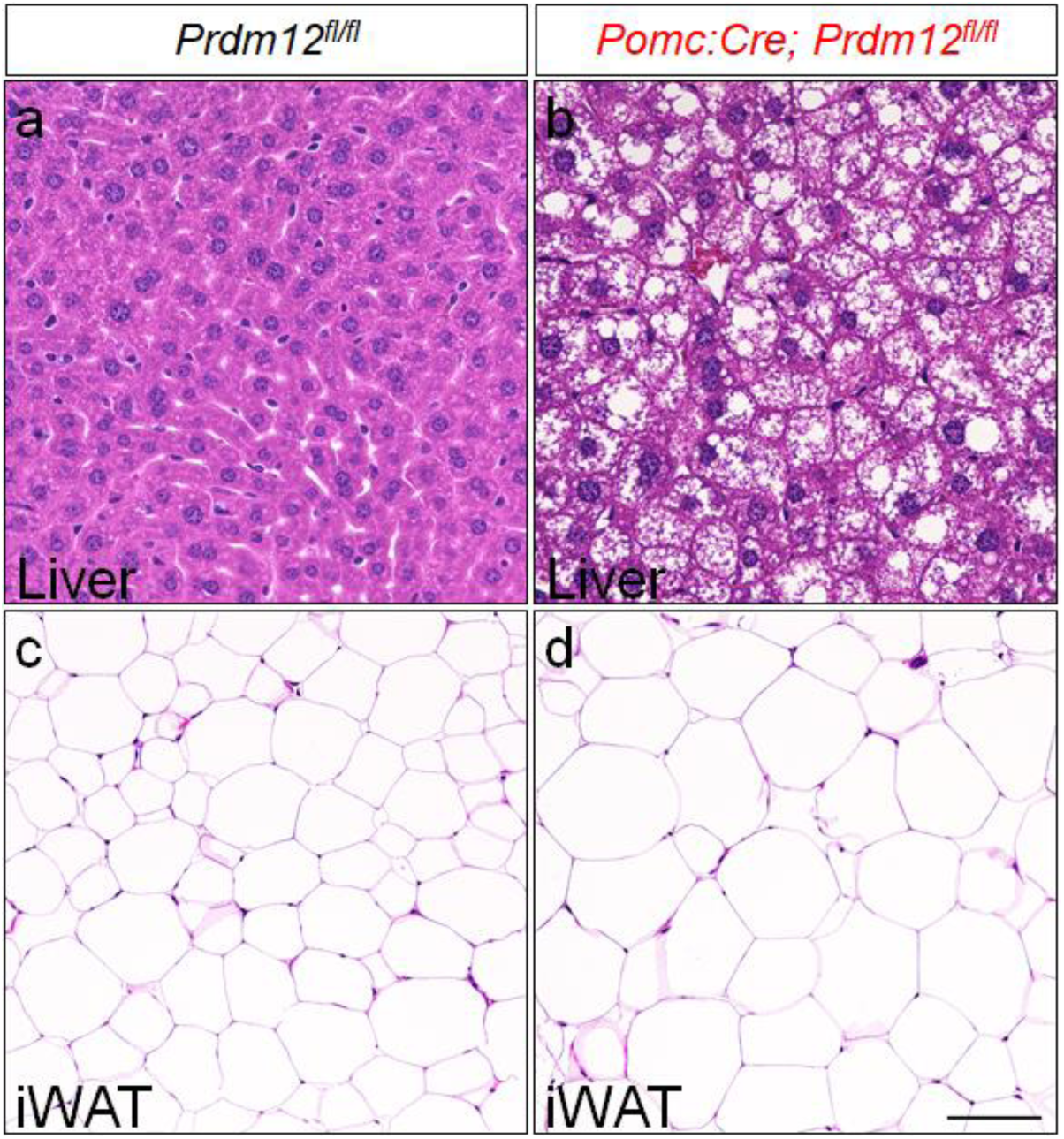
Excessive lipid accumulation in the liver and white adipose tissue of *Prdm12*^*POMCeKO*^ mice. **a, b**, H&E staining in the liver. **c, d**, H&E staining in inguinal white adipose tissue (iWAT). Scale bar is 50 µm for **a, b**, and 100 µm for **c, d**.

## Methods

### Mice

All experimental procedures were approved by the Institutional Animal Care and Use Committee of the UT Southwestern Medical Center. Mice were housed in a barrier facility with a 12-hour light/dark cycle (lights 7:00am-7: 00 pm). Unless otherwise noted, mice were fed standard chow (Envigo 2016 Teklad) and had *ad libitum* access to water.

The *Prdm12*^*fl*^ mice were generated using CRISPR-Cas9 based genome-editing. Two guide RNA sequences (CATGACCCGCTGCGCCCTTG; CAGCAGCTTTTCGTTACCGA) were designed *in silico* using CRISPOR (crispor.tefor.net) flanking the Exon 5. Single-stranded DNA repair templates containing 80 bp homology arms and a *loxP* sequence were synthesized as IDT ultramers. Guides, tracRNA, ssDNA repair template and Cas9 protein (IDT) were mixed by the UTSW Transgenic Core for pronuclear injection. We used the following primers to genotype the *Prdm12*^*fl*^ (248bp) and the *Prdm12*^*+*^ (200bp) alleles: 5’ GGTGCAAGGTCCAAATGTCT 3’ and 5’ CACACTGCAGCTGGCTAAAC 3’. *Prdm12*^*Δ*^ (401bp) was detected using 5’ TTCGGATGAGATGGCTAAGG 3’ and 5’ CCAATGTGCCACAGTACAGG 3’. POMC^eGFP^ (#009593), NPY^hGFP^ (#006417), R26R^Ai14^ (#007908) and POMC-Cre (#010714) mice are available at the Jackson Laboratory.

### Flow cytometry and total RNA isolation

Under a fluorescent dissecting microscope, we extracted the arcuate nucleus from *Pomc*^*eGFP*^ and *Npy*^*hrGFP*^ embryos at E15.5. Cells were disassociated as described previously with minor modifications (Wyler et al 2016 PMID: 26843655). Specially, dissected tissue was collected in solution consisting of Hybernate E (Life Technologies) supplemented with 2% B27 (ThermoFisher) and 0.25% Glutamax (Invitrogen). Tissue was washed with PBS and incubated on a shaker at 37°C for 30 minutes in 20 U/ml papain, 1mM L-Cystine, 0.5mM EDTA, 0.05 mg/ml DNaseI in Earle’s Balanced Salt Solution using the Papain Disassociation System (Worthington). Tissue was then washed 3 times in PBS and resuspended in Hybernate E/B27/Glutamax solution described above. Cells were disassociated with a fire-polished glass pipette. Digested cells were sorted using a Becton Dickinson FACS Aria digital cell sorter with an argon laser (200 mW at 488 nm). Each biological replicate was defined as an independent sort from 5-12 pooled embryos from different days from different litters. Total RNA from approximately 5-10 thousand cells/replicate was isolated using chloroform extraction with 10 µg of Glycoblue (Ambion) used as a carryover, RNA was cleaned and DNase I treated using the RNA Clean & Concentrator-5 kit (Zymo, Irvine CA). RNA quantity and quality were assessed using an Agilent 2100 bioanalyzer. All samples had an RNA integrity number (RIN) ≥9.3. Libraries were prepared using the SMARTer® Stranded Total RNA-Seq Kit v2 (Clonetech, Mountainview, CA) using 1 ng of total RNA and sequenced using an Illumina HiSeq 2500.

### RNA-Seq data analysis

Data were analyzed in R using Bioconductor package. Reads were aligned to mouse reference genome GRCm38/mm10 assembly using HISAT2. We obtained between 71.9 and 87.6 million mapped reads/sample (average of 77.8 million). Duplicates were removed using differential expression analysis performed using Bioconductor version 3.8 packages rSamtools, edgeR version 3.8 (Li et al 2009 PMID 19505943, Robinson et al. 2010 PMID 19910308), using a cutoff of 25 counts per million and a fold-change cutoff of 2 and an FDR of 5%, q-value (Benjamin-Hochberg correction). List of transcription factors was generated by combining databases from GO term (0140110), Riken Institute (genome.gsc.riken.jp/TFdb), and the TFCheckpoint database (www.tfcheckpoint.org) (Tripathi et al, 2013 PMID: 23981286). This yielded a correlated list of 2476 TFs in the mouse genome. Gene ontology (GO) was performed with WebGestalt 2019 (Zhang et al. 2005 PMID: 15980575) using overrepresentation enrichment analysis (OEA) with a minimum of 3 genes per GO category and an FDR of 5% (hypergeometric test, with a BH correction). Data files are deposited in Gene Expression Omnibus (GEO) accession number (xxxxxx)

### Immunohistochemistry, *in situ* hybridization, and qPCR analyses

Details for performing these experiments were described previously (Liu et al., 2010 PMID: 20818386).

The primary antibodies used include anti-GFP (#ab13970, Abcam), anti-Nkx2.1 (#sc-13040, Santa Cruz Biotechnology), anti-Tcf7l2 (#2569, Cell Signaling Technology), anti-ACTH (#AFP-156102789, National Hormone & Peptide Program), and anti-beta-endorphin (#H-022-33, Phoenix pharmaceutical). Secondary antibodies (Alexa Fluor 488 or 594) were from Thermo Fisher.

The following primer sets were used to generated PCR templates for synthesizing digoxigenin (DIG)-labeled RNA probes. Pomc_T3S: 5’ GAATTAACCCTCACTAAAGGGGCTTGCAAACTCGACCTCTC 3’; Pomc_T7AS: 5’ GTAATACGACTCACTATAGGGCTCTTGATGATGGCGTTCTTG 3’; Agrp_T3S: 5’ GAATTAACCCTCACTAAAGGGCCCAAGAATGGACTGAGCA 3’; Agrp_T7AS: 5’ GTAATACGACTCACTATAGGGCGCAAAAGGCATTGAAGAAGC 3’; Npy_T3S: 5’ GAATTAACCCTCACTAAAGGGCCGGTGGATCTCTTCTCTCA 3’; Npy_T7AS: 5’ GTAATACGACTCACTATAGGGCGATGAGGGTGGAAACTTGGA 3’.

RNAscope was performed using ACD RNAscope Multiplex Fluorescent Detection Kit v2 following the manufacturer’s (Advanced Cell Diagnostics) protocol with the following probes Mm-Pomc, Mm-Npy, Mm-Prdm12-C2, and Mm-Nr5a1-C2.

Quantitative PCR (qPCR) was performed using an QuantStudio 6 Flex Real-Time PCR System (Applied Biosystems) and TaqMan Universal PCR Master Mix (4364338). TaqMan probes for *Agrp* (Mm00475829_g1), *Npy* (Mm01410146_m1), *Pomc* (Mm00435874_m1), *Cartpt* (Mm04210469_m1), *Kiss1* (Mm03058560_m1), *Sst* (Mm00436671_m1), *Th* (Mm00447557_m1), *Lepr* (Mm00440181_m1), *Gad1* (Mm04207432_g1), and *Gad2* (Mm00484623_m1) were from ThermoFisher.

### Metabolic analysis

Details for performing body weight, length, body composition, GTT, ITT, blood chemistry, and metabolic chamber analyses were described previously (Liu et al., 2014 PMID:24703703).

### Statistics

Data represented as mean ± s.e.m. Statistical differences were analyzed using paired/unpaired two-tailed Student’s t-test or two-way ANOVA with Sidak’s post hoc testing where applicable (Prism 7.0). Statistical analyses for RNA-seq experiments can be found in **RNA-seq data analysis**. The statistical parameters (i.e. the exact n numbers, p values, and the number of biological repeats) can be found in figure legends.

## Data availability

The data that support the findings of this study are available from the corresponding author upon request. RNA sequencing data reported in this paper have been deposited into NCBI’s Gene Expression Omnibus under accession number GEO XXXXX:

## Acknowledgments

The authors were supported by U.S. NIH grants R01 DK114036 to C.L.; F32DK116427 to S.C.W. C.L. was also supported by an AHA Scientist Development Grant 16SDG27260001, a UTSW Pilot & Feasibility Award, and a Grossman Endowment Award for Diabetes Research. We would like to thank members of the Next Generation Sequencing Core, the Transgenic Core, the Histo Pathology Core, the Flow Cytometry Core, and the Metabolic Phenotyping Core at UTSW.

## Affiliations

Department of Internal Medicine, UT Southwestern Medical Center, Dallas, Texas, USA

Xiameng Chen, Steven C. Wyler, Li Li, Amanda G. Arnold, Rong Wan, Chen Liu

Department of Neuroscience, UT Southwestern Medical Center, Dallas, Texas, USA

Pin Xu, Chen Liu

## Contributions

C.L. conceived the study and generated *Prdm12*^*fl*^ mice. X.C. and S.C.W. performed FACS. S.C.W. and P.X. performed RNA-seq data analysis. X.C., L.L., A.G.A., and R.W., handled all the mouse experiments, performed immunohistochemistry and in situ hybridization. X.C., S.C.W., P.X., and C.L. designed the experiments, analyzed the data, and wrote the manuscript.

## Supplementary information

Supplementary Figures 1-3 and Supplementary Tables 1-4.

